# Ubiquitination and degradation of NF90 by Tim-3 inhibits antiviral innate immunity

**DOI:** 10.1101/2021.01.13.426507

**Authors:** Shuaijie Dou, Guoxian Li, Ge Li, Chunmei Hou, Yang Zheng, Lili Tang, Yang Gao, Rongliang Mo, Yuxiang Li, Renxi Wang, Beifen Shen, Jun Zhang, Gencheng Han

## Abstract

Nuclear Factor 90 (NF90) is a novel virus sensor that serves to initiate antiviral innate immunity by triggering the stress granules (SGs) formation. However, the regulation of the NF90-SGs pathway remain largely unclear. We found that Tim-3, an immune checkpoint inhibitor, promotes the ubiquitination and degradation of NF90 and inhibits NF90-SGs mediated antiviral immunity. Vesicular Stomatitis Virus (VSV) infection induces the up-regulation and activation of Tim-3 in macrophages which in turn recruited the E3 ubiquitin ligase TRIM47 to the zinc finger domain of NF90 and initiated a proteasome-dependent degradation of the NF90 via K48-linked ubiquitination at Lys297. Targeted inactivation of the Tim-3 enhances the NF90 downstream SGs formation by selectively increasing the phosphorylation of PKR and eIF2a, the expression of SGs markers G3BP1 and TIA-1, and protected mice from lethal VSV challenge. These findings provide insights into the crosstalk between Tim-3 and other receptors in antiviral innate immunity and its related clinical significance.

## Introduction

Innate immunity is the first line of host defense against viral infection. Pattern recognition receptors (PRRs), including Toll-like receptors (TLRs) and RIG-I-like receptors (RLRs) are main sensors in defending virus infection(*1*). PRR-mediated downstream signaling pathways initiating an anti-viral innate immune response is the classic anti-virus infection model(*2, 3*). Recently studies have found that nuclear factor 90 (NF90), which is encoded by interleukin enhancer-binding factor-3, is a critical sensor for invading viruses(*4-8*). NF90 is an evolutionarily conserved member of the dsRNA-binding protein family and is abundantly expressed in various mammalian cells(*9, 10*). As an important antiviral pathway, NF90 recognizes virus dsRNA and triggers the formation of stress granules (SGs), which is composed of cytoplasmic particles including ribonucleoproteins, RNA-binding proteins, and translation initiation factors(*11*). Unfortunately, a growing number of virus families modulate SG formation and function to maximize replication efficiency(*12*). Therefore, an understanding of the precise regulation mechanisms of NF90-SGs signaling for efficient viral clearance without harmful immunopathology is needed.

Upon sensing virus, NF90 induces the phosphorylation of double-stranded RNA (dsRNA)-activated kinase protein kinase R (PKR)(*4, 7*). Then SGs form following PKR-mediated phosphorylation and activation of the eukaryotic translation initiation factor 2α (eIF2α), which induces the expression of G3BP1(*13*) (Ras-GAP SH3-binding protein-1) and TIA-1 (T-cell intracellular antigen-1), two key markers in SGs. The activated eIF2α co-operate with other components of SGs to block the virus mRNA translation. Despite acting as an important virus sensor and trigger of SG formation, how NF90 is regulated remains largely unknown.

Tim-3 is an immune checkpoint inhibitor which was first identified in activated T cells. Later Tim-3 was also found to be expressed in innate immune cells, such as dendritic cells and macrophages(*14*). Establishment of Tim-3 as an exhaustion marker in immune cells of both tumors and infectious diseases makes Tim-3 an attractive target for immunotherapy similar to PD-1(*15*), CTLA-4(*16*), and Siglec-G (*17*). Recently, a report showed that increased Tim-3 expression on immune cells in patients with coronavirus disease (COVID-19) is associated with an exhaustion phenotype(*18*). However, Tim-3 does not have an inhibitory motif within its tail(*19*), and the mechanism by which Tim-3 mediates inhibitory signaling remains largely unclear. Kuchroo VK and colleagues showed that CEACAM1 is a heterophilic ligand of Tim-3 and is required for Tim-3 to mediate T cell inhibition(*15*) and that Bat-3 acts as a safety catch, which blocks Tim-3-mediated inhibitory signals in T cells(*17*). Potentiating anti-infection immunity by inducing innate immune responses is a promising area of infection therapy. However, little is known about the Tim-3 signaling in innate immune cells.

Ubiquitination is one of the most versatile posttranslational modifications and is indispensable for antiviral infection(*20*). Increasing evidences suggests that ubiquitination play important roles in various cellular processes, including cell proliferation and antiviral innate signaling. Posttranslational modification of many signaling molecules, including TRAF3/6(*21*), RIG-I(*1*), MAVS(*22*), TBK1(*23*), IRF3/7(*24, 25*), and NLRP3(*26*) involved in TLRs, RLRs, and NLRs pathways by different types of ubiquitination play key roles in the regulation of antiviral innate immunity. However, whether NF90, a molecule containing a ubiquitin binding domain (domain associated with zinc fingers, DZF), undergoes ubiquitination remains unclear.

Here we found that Tim-3 was involved in innate immunity against VSV by promoting the proteasomal degradation of NF90 via the tripartite motif-containing protein 47 (TRIM47)-mediated conjugation of K48-linked ubiquitin. To the best of our knowledge, it is the first time demonstrating the ubiquitination as a new post translational modification mechanism of NF90. Our findings shed a new light on the Tim-3-mediated immune tolerance during infection.

## Results

### Tim-3 interacts with and inhibits NF90

To test whether Tim-3 is involved in the innate immunity against virus, we challenged macrophages with Vesicular Stomatitis Virus, an RNA virus widely used for investigating anti-viral immunity in both mouse and human models *(1)*. Shortly after VSV challenge, the expression of Tim-3 was upregulated in macrophages (Fig.1A&B). The phosphorylation of Tim-3, which account for the Tim-3 signaling(*19*), were also tested. There was a time-dependent enhancement of Tim-3 phosphorylation in HEK293T cells transfected with Tim-3 and challenged with VSV (Fig. 1C). To evaluate the role of the VSV activated Tim-3 in host antiviral innate immune response, macrophages from *Tim-3* knockout mice (*Tim-3*^−/−^ mice) and from wildtype mice (*Tim-3*^+/+^ mice) and the macrophage cell line RAW264.7 cells with a knockdown of Tim-3 (si-*Tim-3*) were challenged with VSV. Both knockout or knockdown Tim-3 in macrophages led to decreased VSV replication (Fig. 1D&E). These findings suggest a negative regulatory role of Tim-3 in anti-viral innate immunity.

**Fig 1.**
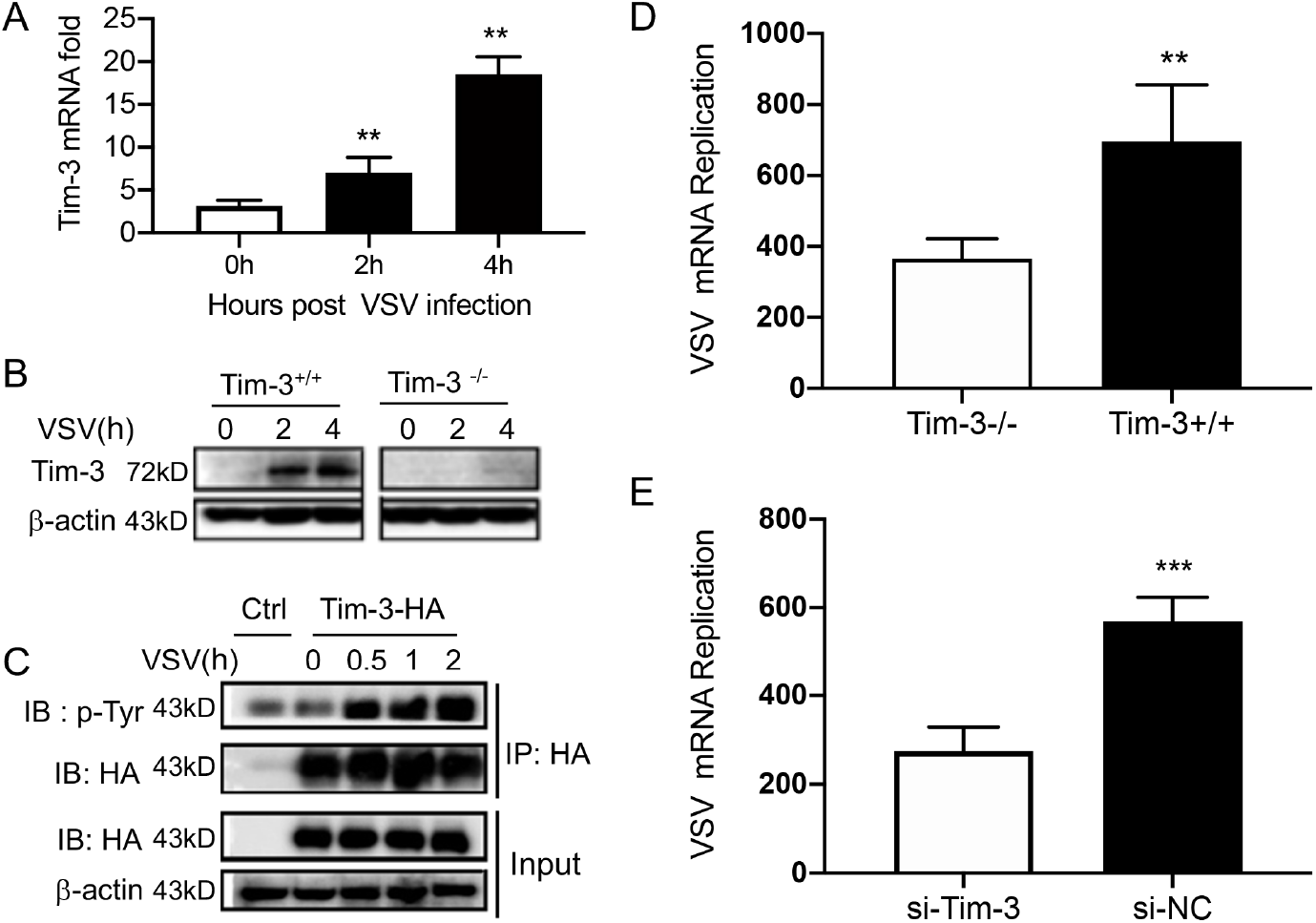
Tim-3 inhibits VSV replication in macrophages. (A) qPCR analysis of Tim-3 mRNA expression in RAW264.7 macrophages infected with VSV for the indicated hours. (B) Peritoneal macrophages were isolated from wild type (Tim-3^+/+^) and Tim-3 knock out mice (Tim-3^−/−^) and were infected with VSV for indicated times. Then the expression of Tim-3 were analyzed by western blot analysis. (C and D) Peritoneal macrophages obtained from Tim-3^+/+^ and Tim-3^−/−^ mice and RAW264.74 macrophages silenced of Tim-3 (si-Tim-3) and RAW264.7 macrophages (si-NC) were challenged by VSV for 8 hours, then cells were harvested for VSV mRNA replication by qPCR. The results shown in all panels were performed three times. **p<0.01, ***p<0.001

To find the possible mechanisms of Tim-3 mediated anti-viral immunity, we used Tim-3 pulldown and mass spectrometry to identify the proteins interacting with Tim-3. Among the candidates, NF90, an RNA-binding protein involved in anti-infection and anti-tumor immunity, received the highest score and the highest number of matched peptides (Supplemental Fig.1). To confirm the interaction between Tim-3 and NF90, HEK293T cells were co-transferred with NF90 and Tim-3, immunoprecipitation targeting either Tim-3 (Fig. 2A) or NF90 (Fig. 2B) all confirmed the interaction between Tim-3 and NF90. The interaction sites of Tim-3 interacting with NF90 were subsequently examined. The results showed that 4Y/F mutant of Tim-3 (Y265F, Y272F, Y280F, Y281F) weakened the binding of Tim-3 with NF90. While deletion of the intracellular domain of Tim-3 (ΔIC) lost the binding activity of Tim-3 with NF90 (Fig 2C&D). These data showed that Tim-3 interacts with NF90 through its intracellular domain, in which Y265, Y272, Y280, and Y281 play an important role. We finally evaluated the effects of Tim-3 on NF90 expression. In HEK293T cells transfected with Tim-3 or in macrophages from Tim-3 transgenic mice (Tim-3-TG), the overexpression of Tim-3 decreased the expression of NF90 (Fig 2E). These results show that Tim-3 interacts with and inhibits NF90.

**Figure 2.**
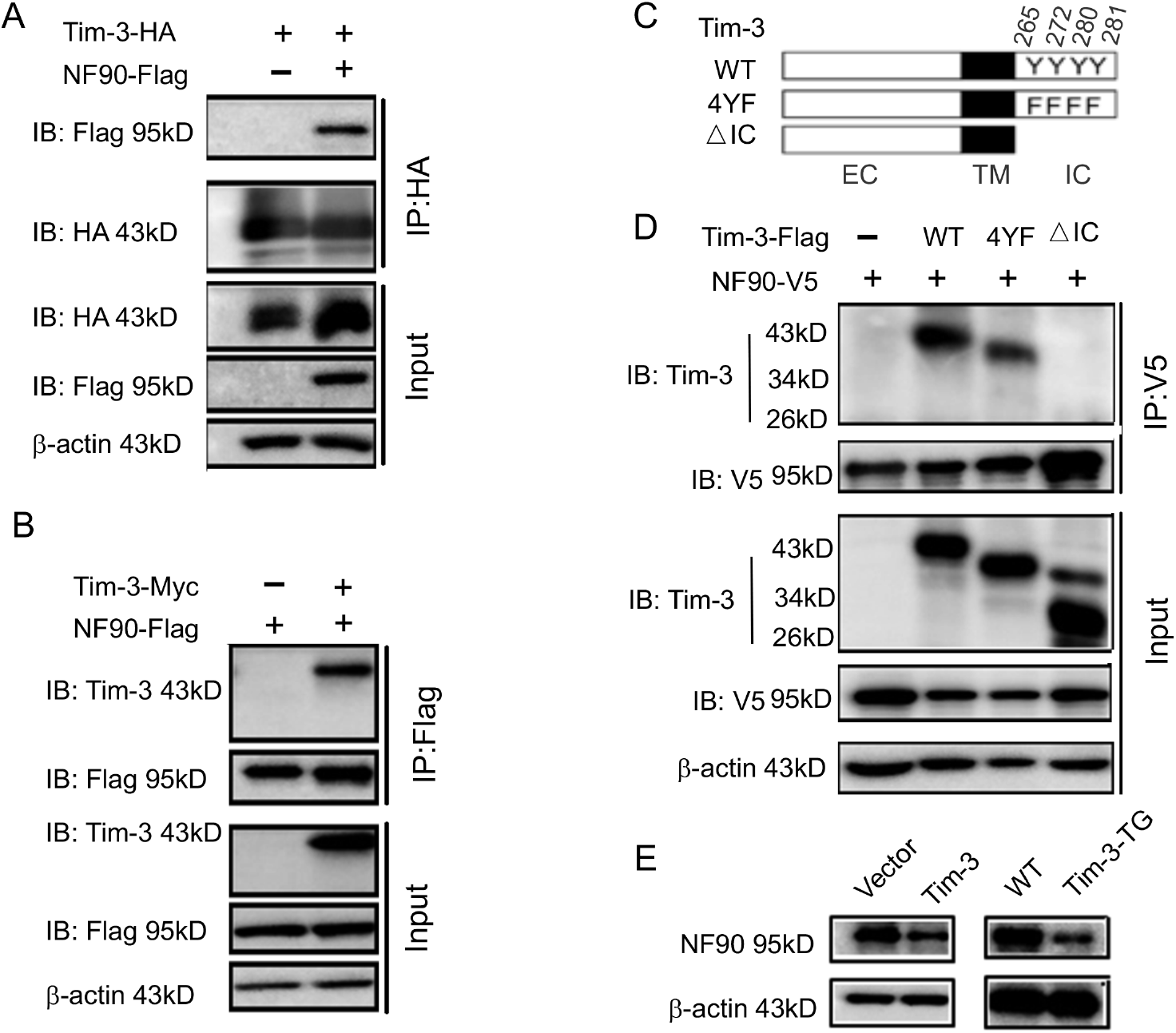
Tim-3 interacts with and inhibits NF90. (A and B) Protein complex of Tim-3 and NF90 overexpressed in cells. HEK293T cells were transfected with plasmids encoding HA-Tim-3, Flag-NF90, and Myc-Tim-3 for 24 h, immunoprecipitated with HA or Flag antibody, respectively, and detected by western blot for the indicated antibodies. (C and D) Interaction of Tim-3 intracellular domain with NF90. Schematic structure of Tim-3 and the derivatives used are shown (C). Whole cell lysis of HEK293T cells transfected with Flag-Tim-3 (WT), Flag-Tim-3 (Del), Flag-Tim-3 (4YF), and V5-NF90 were used for immunoprecipitation and immunoblotting, as indicated (D). (E) Immunoblot analysis of NF90. HEK293T cells were transfected with Tim-3 plasmid for 24 h, and lysates were detected for NF90 expression by western blot (left). Peritoneal macrophages from WT and Tim-3-TG mice were lysed and NF90 protein were detected by western blot (right). The results shown in A, B, D,F were performed three times.

### Tim-3 promotes the ubiquitination of NF90 at the DZF domain

Ubiquitination is one of the most versatile posttranslational modifications and is indispensable for antiviral immunity. However, whether NF90 undergoes proteasomal degradation is totally unknown. To find the mechanisms by which Tim-3 inhibits NF90 expression, we tested whether NF90 undergoes ubiquitination and if so, whether Tim-3 is involved. Macrophages isolated from wild type (WT) and Tim-3 transgenic mice (Tim-3-TG) were challenged by VSV for 4 hours and the ubiquitination of NF90 were examined. Interestingly, the ubiquitination of NF90 were significantly increased in macrophages from Tim-3-TG mice compare to those from wild type mice (Fig. 3A). When NF90, K48-Ub, or Tim-3 was transfected into HEK293T cells, NF90 underwent K48-linked ubiquitination, which can be enhanced by co-transfected Tim-3 (Fig. 3B). Following VSV challenge, the enhanced K48-linked ubiquitination of NF90 was also found in primary macrophages from Tim-3 transgenic mice compared to those from wild type mice (data not shown). The results suggested that Tim-3 may inhibit NF90 by enhancing the K48-linked ubiquitination and degradation of NF90. We then explored the domain of NF90 for ubiquitination using constructions encoding the DZF domain (domain associated with zinc fingers), full-length NF90 or NF90 lack of DZF domain (ΔDZF) (Fig.3C) and found that the DZF domain of NF90 was dominantly ubiquitinated (Fig. 3D). Finally, when the plasmid of Tim-3 was co-transfected with different NF90 constructs, the data revealed that Tim-3 specially enhanced the ubiquitination of DZF (Fig. 3E). These results demonstrated that NF90 can be ubiquitinated at the DZF domain, and the process can be enhanced by Tim-3, suggesting that Tim-3 may suppress NF90 by promoting its ubiquitination and degradation.

**Figure 3.**
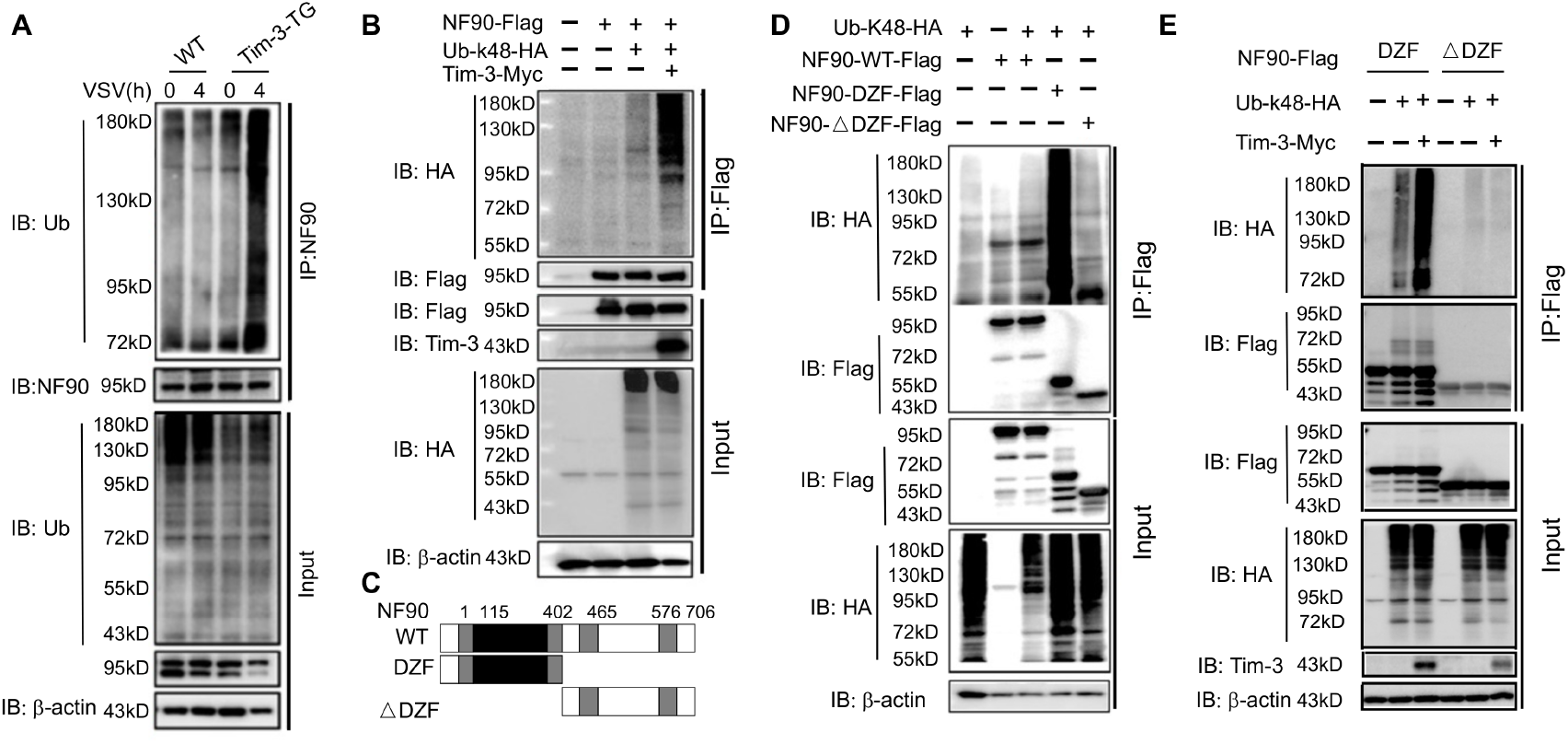
Tim-3 promotes the ubiquitination of NF90 at the DZF domain. (A) Tim-3 enhances the ubiquitination of NF90 in macrophages in response to VSV challenges. Peritoneal macrophages in WT and Tim-3-TG mice were infected with VSV for 4 h and cells were treated with MG132 (20 ug/ml) for 6 h before harvesting protein lysates, followed by western blot analysis of the total-Ub of NF90 immuno-precipitated with antibody to ILF3 (NF90). (B) Tim-3 promotes the ubiquitination of NF90. HEK293T cells were transfected with plasmids encoding Flag-NF90 or HA-Ub-K48, Tim-3-Myc for 24 h, treated with MG132 (20 ug/ml) for 6 h, immunoprecipitated with Flag antibody, and then detected by western blot for the indicated antibodies. (C) Schematic structure of NF90 and the derivatives used were shown. (D-E) Tim-3 promotes the K48-Ub modification of NF90 at the DZF domain. HEK293T cells were transfected with the indicated plasmids for 24 h and treated with MG132 (20 ug/ml) for 6 h. The cells were then lysed, immunoprecipitated with Flag antibody, and detected by western blot using the indicated antibodies. At least three independent experiments were conducted for all panels.

### Involvement of TRIM-47 in Tim-3 -mediated degradation of NF90

To find the possible E3 ligase accounting for Tim-3 mediated NF90 ubiquitination, NF90-interacting proteins were investigated by immunoprecipitating NF90 and then performing mass spectrometry. Among the NF90-interacting protein candidates, we identified three proteins with potential E3 ligases activities. TRIM47 had the highest Mascot scores and the highest number of matched peptides (Fig. 4A). Knockdown of TRIM47 with specific siRNA (si-TRIM47) in macrophages increased the half-life of endogenous NF90 protein during VSV infection (Supplement Fig. 2). To confirm whether the TRIM47 promotes the proteasomal degradation of NF90, we transfected HEK293T cells with ubiquitin, NF90, and an increasing dose of TRIM47. TRIM47 induced degradation of NF90 in a dose-dependent manner which can be blockaded in the presence of MG132, indicating a proteasomal dependent degradation (Fig. 4B). In addition. when *Tim-3* was co-transfected, it dose-dependently enhanced TRIM47-mediated degradation of NF90 (Fig. 4C).

**Figure 4:**
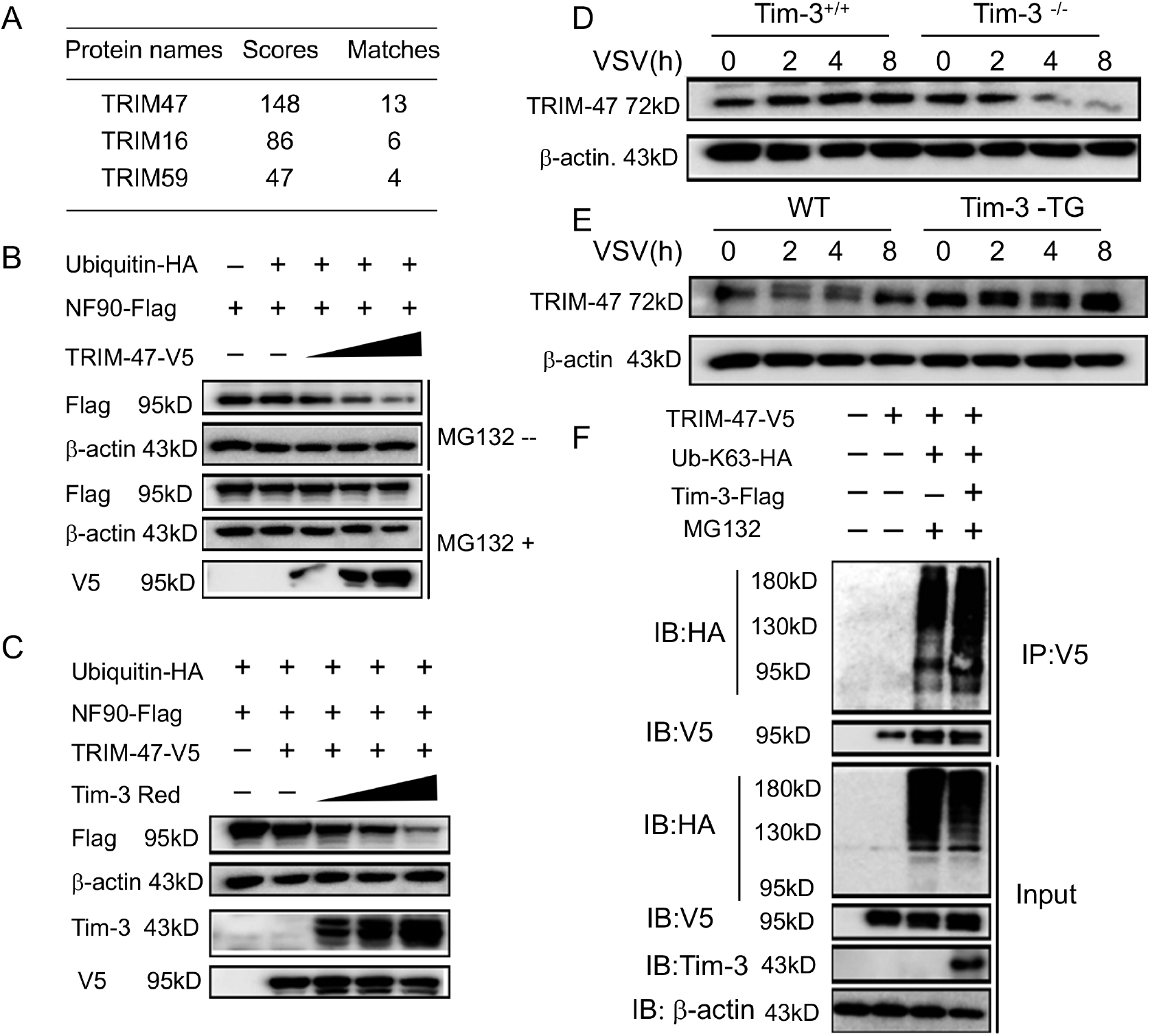
Involvement of TRIM-47 in Tim-3-mediated NF90 degradation. (A) E3 ligases identified by mass spectrometry for top peptide hits (defined by Mascot score) associated with NF90 ubiquitination. (B) TRIM47 promotes NF90 degradation in a proteasome-dependent manner. Plasmids encoding Flag-NF90, HA-Ubiquitin, along with increasing amounts of V5-TRIM47 (0.5, 1.0, and 2.0 ug) were transfected into 293T cells for 24 h, cells were treated with and without MG132 (20 ug /ml), respectively, followed by western blot to examine the NF90 protein level. (C) Tim-3 accelerates TRIM47-mediated NF90 degradation in a dose-dependent manner. HEK293T cells were transfected with plasmids encoding Flag-NF90, HA-ubiquitin and V5-TRIM47 and an increasing dose of plasmid encoding Red-Tim-3 (0.5, 1.0, and 2 ug) for 24 h. The protein level of NF90 was examined in cells. (D and E) Tim-3 upregulates TRIM47 in protein levels.TRIM47 protein levels were analyzed by Immunoblot in lysates from *Tim-3*^*+/+*^ or *Tim-3*^−/−^ and WT or Tim-3-TG macrophages infected with VSV for the indicated time. (F) Tim-3 facilitates K63-linked ubiquitination mediated by TRIM47. Plasmids encoding HA-Ub-K63, V5-TRIM47, and Flag-Tim-3 were transfected into HEK293T cells. Cells were treated with MG132 (20 ug/ml) for 6 h, and cell lysates were immunoprecipitated with Flag antibody and detected by western blot for K48-Ub levels. At least three independent experiments were conducted for all panels.

We then explored the possible interaction between Tim-3 and TRIM47. VSV challenge led to decreased TRIM47 expression in *Tim-3*^−/−^ cells compared with that in *Tim-3*^*+/+*^ cells (Fig. 4D), and increased TRIM47 expression in Tim-3 transgenic mice-derived macrophages compared with that in control cells (Fig. 4E). These results showed that Tim-3 promotes the expression of TRIM47 in the presence of virus. The possible mechanism by which Tim-3 enhances TRIM-47 expression was primarily investigated. We examined whether TRIM-47 undergoes ubiquitination when Tim-3 is overexpressed, as TRIM25, an E3 ligase with a structure similar to TRIM-47, undergoes ubiquitination during viral infection(*27, 28*). Interestingly, when the genes encoding TRIM47, Tim-3, and Ub-K63 were co-transfected into HEK-293T cells, TRIM47 underwent ubiquitination, and Tim-3 enhanced this progress (Fig. 4F). However, the relationship between Tim-3 enhanced TRIM47 expression and Tim-3 enhanced TRIM47 ubiquitination remains to be determined. These results showed the involvement of TRIM47 in Tim-3 mediated degradation of NF90.

### Tim-3 recruits TRIM-47 to the DZF domain of NF90 and Lys297 within DZF is a Critical Site for TRIM47-Mediated K48-Linked Ubiquitination of NF90

To find whether Tim-3 and TRIM-47 interacts with each other to act on NF90, we first examined the interactions among Tim-3, TRIM47, and NF90. Different Tim-3 constructs (Fig.5A) or different NF90 constructs (Fig.5B) were co-transfected with TRIM-47 into HEK293T cells. The immunoprecipitation assay showed that TRIM-47 interacted with the intracellular domain of Tim-3, and interacts with the DZF domain of NF90. These data suggest that Tim-3 recruits TRIM47 to the DZF domain via its intracellular domain where forming a complex of TRIM47 and NF90.

To confirm that Tim-3 co-operates with TRIM47 to enhance the ubiquitination and degradation of NF90, we co-transfected Tim-3, TRIM47, and ubiquitin into HEK293T cells and examined the effects of TRIM47 and Tim-3 on the ubiquitination of NF90. Overexpression of TRIM47 promotes the ubiquitination of NF90, and that this process was enhanced when Tim-3 was co-transfected (Fig. 5C). Next, we examined the ubiquitin modification site within the DZF domain of NF90 by site mutation. NF90 contains eighteen Lysine residues in its DZF domain. Immunoprecipitation analysis revealed that TRIM47 enhanced the ubiquitination of wildtype DZF and DZF with the K100R, K117/119R, K127R, K143R, K158R, and K224R but not K297R mutants in HEK293T cells (Supplemental Fig.3). We further demonstrated that only mutations of the arginine at K297R completely blocked the TRIM47-mediated ubiquitination and degradation of NF90 via a K48-mediated linkage (Fig. 5D). In addition, when co-transfected K48-Ub, Tim-3, wildtype NF90, DZF domain, or DZF K297R with increased doses of TRIM47, we found that TRIM47 dose-dependently induced the degradation of NF90 DZF domain but not for DZF K297R (Fig. 5E). Taken together, these data suggest that K297 is the critical residue for the ubiquitination of NF90-DZF targeted by TRIM47.

**Figure 5.**
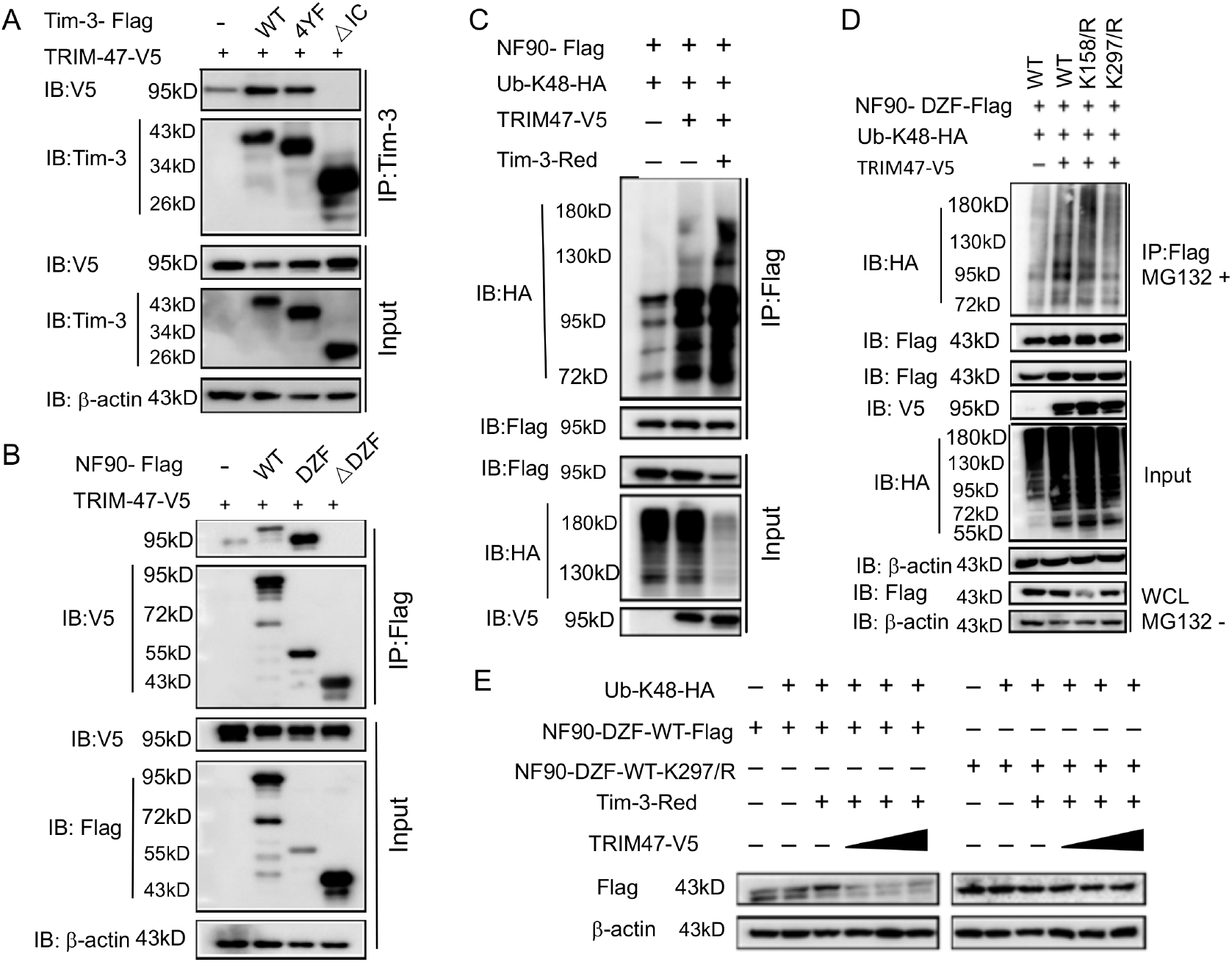
Tim-3 recruits TRIM47 to DZF domain of the NF90 within which Lys297 is a critical site for TRIM47-mediated K48-linked ubiquitination and degradation of NF90. (A&B) The intracellular domain of Tim-3 and the DZF domain of NF90 interacts with TRIM47 respectively. HEK293T cells were transfected with the indicated plasmids for 24 h and treated with MG132 (20 ug/ml) for 6 h. The cells were then lysed, immunoprecipitated with Tim-3 or Flag antibody, and detected by western blot using the indicated antibodies. (C) Tim-3 promotes NF90 degradation mediated by E3 ligase TRIM47. HEK293T cells were transfected with plasmids encoding HA-Ub-K48, V5-TRIM47, Flag-NF90, and Red-Tim-3, and treated with MG132 (20ug/ml) for 6h. Cell lysates were then immunoprecipitated with Flag antibody and analyzed by Western Blot using the indicated antibodies. (D) Residue K297 of NF90 is the major site of TRIM47-mediated K48-linked ubiquitination. Flag-NF90-DZF (WT), or K R mutants, HA-Ub-K48, and V5-TRIM47 were transfected into HEK293T cells for 24 h. Cells were then treated with MG132 (20ug /ml) for 6 h. Cell lysates were analyzed by western blot for K48-linked ubiquitination of NF90. (E) Residue K297 is the decisive site in TRIM47-mediated degradation of NF90. Plasmids encoding Flag-NF90-DZF (WT), or K297R mutants, PDsRed-Tim-3, HA-Ub-K48, and V5-TRIM47 were transfected into 293T cells, and cell lysates were examined by western blot for the indicated proteins. At least three independent experiments were conducted for all panels.

### *Tim-3* Deficiency Enhances the Formation of SGs and Protects Mice from VSV Infection

Finally, the significance of Tim-3 inhibits NF90 was investigated. As NF90 triggers the formation of SGs, we first examined whether Tim-3 regulates the down-stream of NF90-SGs pathway. Peritoneal macrophages were isolated from wildtype and Tim-3 knock out (*Tim-3*^−/−^) mice and following VSV challenge for 2-8 hours, the expression of NF90 and the phosphorylation of PKR, eIF2a, as well as the phosphorylation of other signaling cascade including ERK and P38 were examined. The data in Fig.6A showed that the expression of NF90 and the phosphorylation of eIF2α and PKR was dramatically increased in macrophages from *Tim-3*^−/−^ mice compared with that in cells from *Tim-3*^*+/+*^ mice. There was no difference in p38 and ERK phosphorylation between *Tim-3*^−/−^ and *Tim-3*^*+/+*^ cells. Meanwhile, the expression of G3BP1 and TIA-1, two markers of SGs were also examined in above macrophages. The results in Fig.6B showed that Tim-3 knock out significantly increased the expression of G3BP1 and TIA-1 in macrophages following VSV challenges in vitro. To test the effects of Tim-3 inhibition on NF90-SGS pathway in vivo, a VSV infection model were established in mice. We found that the expression of SG markers: G3BP1 and TIA-1, and VSV replication were significantly higher in spleen, lung, and peritoneal macrophages from *Tim-3*^−/−^ mice than those in *Tim-3*^*+/+*^ mice (Fig 7A-7I). And lethal VSV infections lead to an increased survival rate in *Tim-3*^−/−^ mice compared to *Tim-3*^*+/+*^ mice (Fig. 7J) and a less severe tissue inflammation (Fig. 7K). These data showed that Tim-3 deficiency enhances the formation of SGs in vivo and protects mice from VSV. Finally, we also examined whether the silence of TRIM-47 affects the assembly of SGs and the anti-viral immunity of macrophages. The data showed that silence of TRIM-47 with specific siRNA in RAW264.7 led to increased G3BP1 and TIA-1 expression and decreased virus load following VSV infection (Supplement Fig. 4), further confirming that TRIM47 acts as an up-stream regulator of the NF90-SG antiviral pathway.

**Figure 6:**
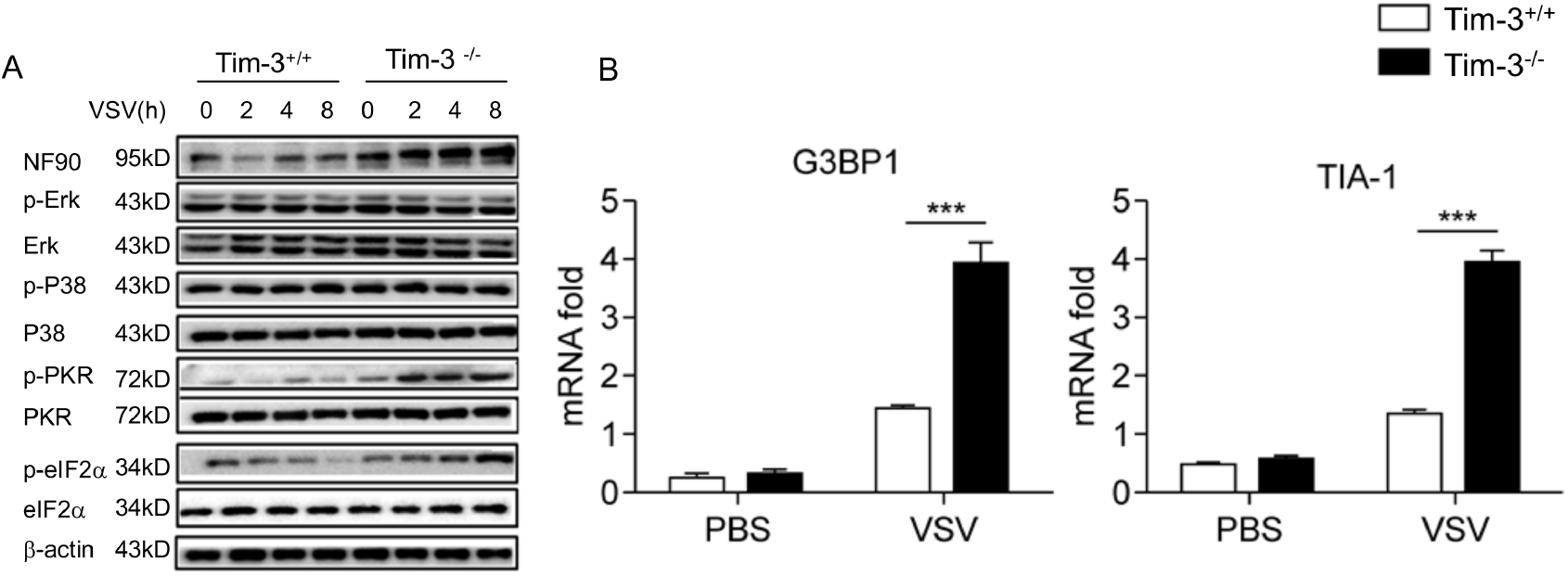
Tim-3 selectively inhibits the phosphorylation of PKR and eIF2a, increases the expression of SGs markers G3BP1 and TIA-1 in macrophages. (A) Lysates from *Tim-3*^*+/+*^ and *Tim-3*^−/−^ peritoneal macrophages infected with VSV were analyzed by immunoblot for the indicated proteins. (B) *Tim-3*^*+/+*^ and *Tim-3*^−/−^ peritoneal macrophages were infected with VSV for 8 hours, and then TIA-1 or G3BP-1 mRNA transcription were analyzed by qPCR. The results shown are representative of three independent experiments. ***,p<0.001

**Figure 7.**
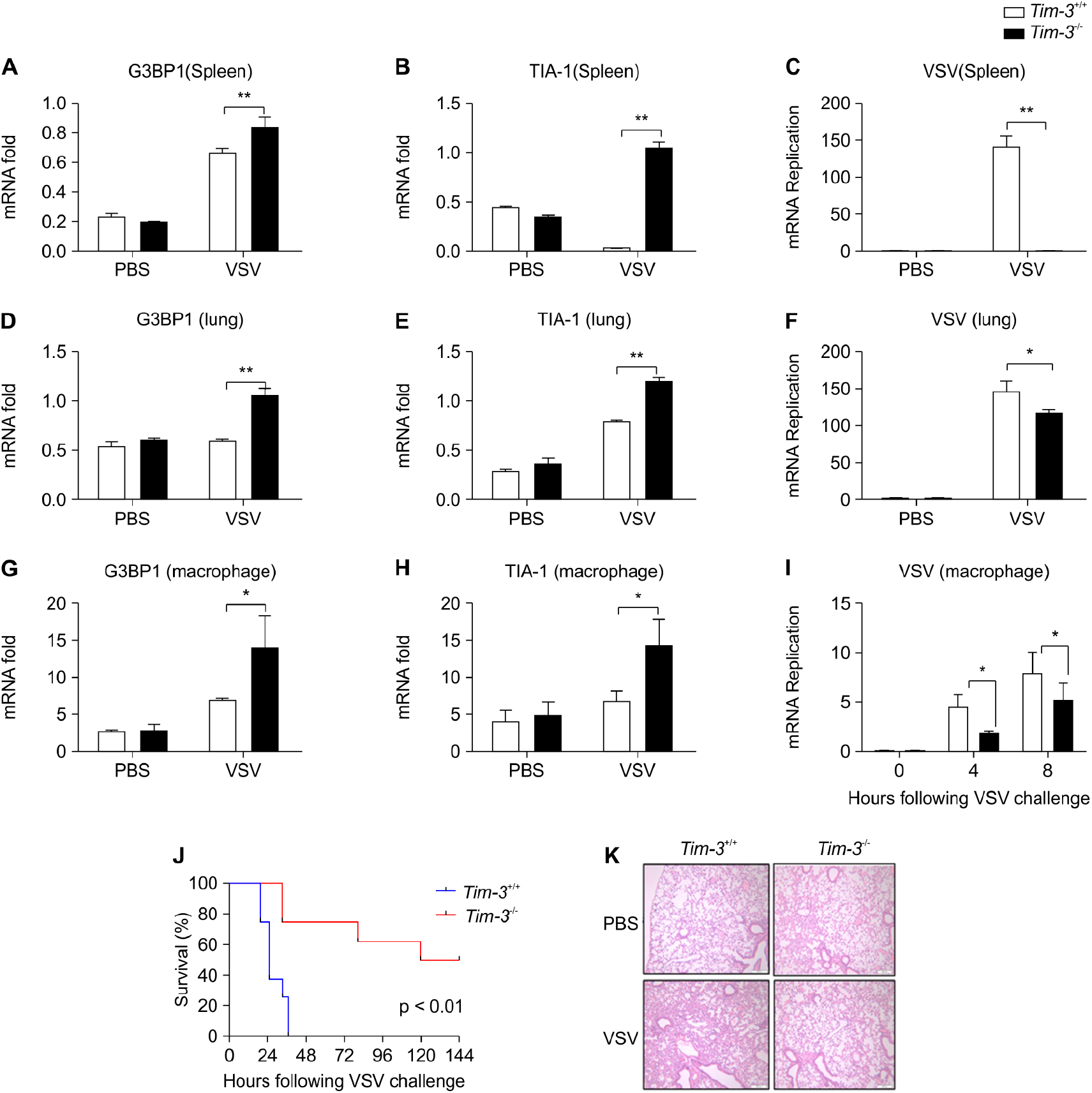
Tim-3 deficiency upregulates G3BP1 and TIA-1 and protects mice from VSV infection. **(A**, B and D, E and G, H) Detection of mRNA transcription of G3BP1 and TIA-1 in organs and peritoneal macrophages by qPCR after *Tim-3+/+* and *Tim-3*^*-/ -*^mice (n = 5 per group) were intraperitoneally injected with VSV for 24 h. (C, F and I) qPCR analysis of VSV loads in organs and peritoneal macrophages after *Tim-3*^*+/+*^ and *Tim-3-/-* mice (n = 5 per group) were intraperitoneally injected with VSV for 24 h. (J) *Tim-3*^*+/+*^ and *Tim-3*^−/−^ mice (∼7-week-old) were intraperitoneally injected with VSV (1× 10^8^ pfu/g) (n = 10 per group) followed by recording survival of both groups. *P* < 0.01. (K) The lung tissues from *Tim-3*^*+/+*^ and Tim-3^−/−^ mice (C, F, and I) were stained with hematoxylin and eosin, and their pathology analyzed in response to VSV. The results shown are representative of three independent experiments. *, p<0.05; **p<0.01.

The mechanisms by which Tim-3 promotes the TRIM47-mediated ubiquitination and proteasomal degradation of NF90 in viral immune evasion are summarized in Fig. 8. Upon VSV infection, Tim-3 is activated and upregulated. The activation of Tim-3 in turn recruited the E3 ubiquitin ligase TRIM47 to the zinc finger domain of NF90 and initiated the proteasome-dependent degradation of the NF90 via K48-linked ubiquitination at Lys297. The negative regulation of NF90 by Tim-3 blocked the virus triggered and NF-90-SGs mediated antiviral immunity, and finally led to virus immune evasion.

**Figure 8:**
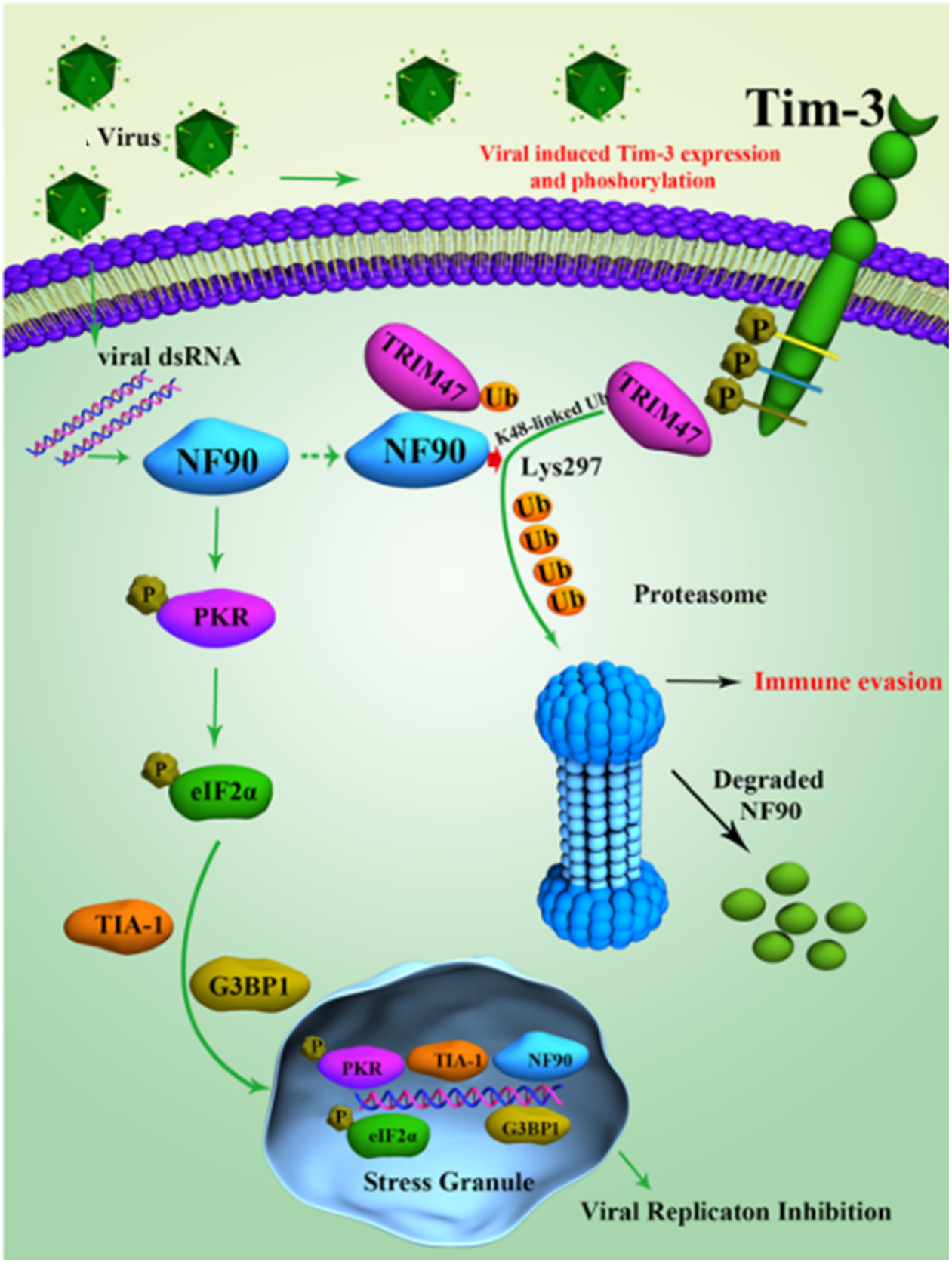
Schematic diagram of how Tim-3 inhibits NF90-SGs pathway in macrophages during infection. Upon VSV infection, Tim-3 is activated and upregulated. The activation of Tim-3 in turn recruited the E3 ubiquitin ligase TRIM47 to the zinc finger domain of NF90 and initiated a proteasome-dependent degradation of the NF90 via K48-linked ubiquitination at Lys297. The negative regulation of NF90 by Tim-3 blocked the RNA virus triggered and NF-90-SGs mediated antiviral immunity, and finally led to virus immune evasion.

## Discussion

Viruses have developed many different strategies of immune evasion, for example by downregulating or degrading virus sensors. The mechanisms by which these receptors are regulated are widely investigated in hopes of developing effective treatments. NF90 was found to play an important role in host innate immunity against various virus infections However, the regulation of NF90 under physio-pathological conditions remains largely unclear. Here we identified a novel negative regulation mechanism of NF90, which could be employed by VSV to evade the immune attack. The VSV activated Tim-3 in turn promotes the ubiquitination and degradation of NF90, and subsequently inhibits the formation and the antiviral activity of down-stream SGs. To our best knowledge, this is the first report showing that NF90 undergoes ubiquitination and also the critical domain (DZF) and critical site (Lys297) of NF90 for Tim-3 and E3 ligase TRIM47 mediated ubiquitination and degradation.

NF90, like the classical PRRs, is considered a novel virus sensor, exerting an important role in host innate immunity against various viral infections, especially negative-sense single-stranded RNA virus(*4, 6, 7*). One report showed that NF90 is required for an efficient response against VSV infection(*8*), but the underlying have not been clarified. Other report showed that NF90 interacted with the VP35 protein of Ebola virus and inhibited EBOV infection through impairing the function of the EBOV transcription/replication complex(*5*). Knockdown of NF90 in indicated cells dramatically promote EBOV and influenza virus replication, while overexpression of NF90 inhibits or impacts replication of these viruses(*4-6*). When the signaling cascades are investigated, SGs, not the interferon pathway, serve as the downstream signaling cascade of the NF90 antiviral pathway (*5,8*). NF90 promotes(*29*) the assemblage of SG and synergizes with other proteins to exert antiviral immunity(*7*). These reports support our findings that NF90 inhibits VSV replication via SGs and further demonstrates the critical role of the NF90-SG signaling pathway during an antiviral immune response.

Tim-3 is an immune checkpoint inhibitor that was initially found to be expressed on activated Th1 cells by binding with its ligand Gal-9(*30, 31*).Tim-3 induces apoptosis and T cell tolerance(*32, 33*). Most investigations focus on the roles of Tim-3 in maintaining T cell exhaustion in immune disorders, tumors, or infectious diseases, which means that this checkpoint inhibitor could be abused. Recently our works and other published data focus on the roles of Tim-3 in maintaining the homeostasis of innate immune cells and demonstrated that the dysregulated Tim-3 on innate immune cells also contribute to the immune evasion of many tumors or pathogens(*18, 34*). Innate immune cells have now attached much attention for developing new therapeutic strategies against infectious or tumors diseases, in which innate immune cells expressed immune checkpoint inhibitors are potential targets (*14*). Here we found that Tim-3 suppresses the macrophage-mediated antiviral immune response, suggesting that it is a potent therapeutic target for re-boosting innate immunity against virus. However, as Tim-3 does not possess an obviously classical ITIM motif compared with other immune-checkpoint receptors, such as PD-1, CTLA-4(*35*), and Siglec-G(*1*), the mechanism in which Tim-3 mediates inhibitory signals remain unclear. Kuchroo and colleagues(*15*) showed that Tim-3 induces T cell exhaustion via BAT and using CEACAM1 as a coreceptor, and our findings showed that Tim-3 promotes tumor-prone macrophage polarization by binding to and suppressing the phosphorylation and nuclear translocation of STAT1(*34*). Further, Tim-3 inhibits TLR4-mediated macrophage activation during sepsis by suppressing NF-kB activation(*36*). How Tim-3 works during anti-viral innate immunity remain unclear. The intracellular tail of Tim-3 contains a highly conserved tyrosine-containing src homology 2 (SH2)-binding motif, and tyrosine residues within this motif can be phosphorylated, which is critical for Tim-3 signaling in T cells(*1*). In this study, we identified an increased tyrosine phosphorylation of Tim-3 in macrophages following VSV infection. We also demonstrated that deletion of Tim-3 tail or mutation of the conserved tyrosines, including Tyr265, Tyr272, Tyr280, and Try281, within Tim-3 tail significantly attenuated the interaction between Tim-3 and NF90. These data demonstrated a new mechanism by which Tim-3 transduces inhibitory signal in anti-viral innate immune responses.

Ubiquitination is one of the most versatile posttranslational modifications for substrates and is indispensable for antiviral infection(*37*) However, whether NF90 undergoes ubiquitination is totally unknown. Structurally, NF90 possess a domain associated with zinc fingers (DZF) in the N-terminal region, which is a symbol of ubiquitination for substrates(*38*). Here we verified our hypothesis and demonstrated that NF90 can be ubiquitinated, a process that is enhanced by Tim-3 in macrophages following VSV infection. As speculated, Tim-3 mainly promotes the ubiquitination of the DZF domain of NF90. To our best knowledge, this is the first report demonstrating that NF90 undergoes ubiquitination. Further, we also found that Lys297, a highly conserved residue among different isoforms of NF90, plays a critical role in the K-48 linked ubiquitination of NF90.

When the candidates accounting for NF90 ubiquitination were examined, we focus on TRIM47 as it got the highest scores and the highest number of matched peptides among NF90-interacting proteins, and structurally, TRIM47 contains a RING finger domain in the N-terminus, which may contribute to ubiquitin modification(*38*). In addition, TRIM family proteins play important roles in many biological processes, such as cell cycle regulation, and viral response(*25*). As discussed above, the conserved tyrosines within the Tim-3 tail could form a SH2 (SRC homology 2) binding domain(*17, 19, 34, 39*). We posit that Tim-3 may recruit TRIM47 using its SH2 binding domain as the tyrosines are phosphorylated following VSV infection. A recent report showing that Siglec-G triggers downstream signaling by recruiting SHP-2(*1*) supports this hypothesis. Interestingly, our data showed that Tim-3 enhances the expression and ubiquitination of TRIM47. This is also the first report showing the ubiquitination modification of TRIM47. However, the mechanisms of Tim-3 enhanced TRIM47 ubiquitination and whether the enhanced ubiquitination of TRIM47 by Tim-3 accounts for the increased TRIM47 expression remain to be determined. In summary, we have verified that Tim-3 is specifically upregulated following VSV infection and inhibits NF90 signaling pathway in macrophages. Intracellularly, Tim-3 promotes TRIM47 mediated ubiquitination and degradation of NF90. In VSV infected models, Tim-3 signaling inhibits the formation and the activity of the NF90 downstream SGs. To the best of our knowledge, this is the first report demonstrating the ubiquitination modification of NF90. These findings provide a novel mechanism of Tim-3-mediated infection tolerance which with implication in antiviral applications.

## Materials and Methods

### Mice

The Tim-3-TG mice were produced and fed as described previously(*34*). The Tim-3-flox/flox(*Tim-3*^*+/+*^)mice (C57BL/6) used in this study were generated in the Transgenic Core Facility of Cyagen Biosciences Inc., Guangzhou, China. EII-cre knock-in mice (C57BL/6) were a gift from Dr. Haitao Wu (Institute of Military cognition and Brain Sciences, Beijing, China). *Tim-3*^−/−^(Tim-3^fl/fl^ EII^cre/+^) mice (C57BL/6) were generated by mating Tim-3-flox with EII-cre mice. And the genotype of *Tim-3*^−/−^ *mice* (Havcr2) was detected with a primer set as mHavcr2-Forward: 5’-CCAATTGGGTTCTACTATAAAGCCTTG-3’ and mHavcr2-Reverse1: 5’-AAGTTGAGAGTTCTGGGATTACAGG-3’ and mHavcr2-Reverse2: 5’-ATACTTGCTTCAGTGGCTCGCGA-3’ (Supplemental Fig.S5). Wildtype C57BL/6 mice and the aforementioned Tim-3-TG and *Tim-3*^−/−^ mice were bred in specific pathogen-free conditions. The protocol was approved by the Ethics Committee of Animal Experiments of the Beijing Institute of Brain Sciences. All efforts were made to minimize suffering. Major procedures were blinded.

### Cells and Reagents

The RAW264.7 and HEK293T cell lines (ATCC) were maintained in DMEM (Gibco) supplemented with 10% FBS (Gibco). Peritoneal macrophages were prepared as described(*40*). For stable transfection NC shRNA or Tim-3 shRNA, RAW264.7 macrophages were transfected with Lipofectamine 2000 reagents (Invitrogen, 11668019) and then selected with 1,000 ng/mL G418 (Invitrogen, 10131027), which were purchased from Invitrogen. Vesicular stomatitis virus (VSV) was obtained and cultivated as described(*1, 34*). MG132 was purchased from Selleckchem (S2619) and used at a final concentration of 20 μM.

### Plasmids and Antibodies

The flag-tagged full-length NF90, NF90 mutation plasmids (NF90-DZF and NF90-ΔDZF), V5-tagged full-length NF90, as well as HA-tagged ubiquitin were constructed into eukaryotic pcDNA3.1 (+) -Flag and pcDNA3.1 (+) -V5 eukaryotic expression vector, respectively. Recombinant vectors encoding WT or mutant human-specific Tim-3 were constructed by PCR-based amplification of cDNA from human U937 cells and then subcloned into the eukaryotic pcDNA3.1 (+) eukaryotic expression vectors, with Flag, Myc and HA tags, respectively. Full-length NF90-GFP and full-length Tim-3-RFP fluorescence plasmids were cloned into PEGFP1-N1 and PDsRed1-N1 eukaryotic expression vectors, respectively. TRIM47 full-length for V5-tag plasmids were constructed into the pcDNA3.1 (+) -V5 eukaryotic expression vector. Antibodies to Tim-3 (D3M9R, mouse-specific), Tim-3 (D5D5R, human-specific), eIF2α (D7D3), p-eIF2α (D9G8), Ub-K48 (D9D5), P38 (8690), p-P38 (9215), ERK (4348), and p-ERK (8544) were obtained from Cell Signaling Technology.

An antibody to p-PKR (GTX32348) was purchased from GeneTex. Antibodies to β-Actin, anti-Flag-Tag (CW0287), anti-HA-Tag (CW0092), anti-V5-Tag (CW0094), and anti-RFP (CW0253) were from purchased from CWBIO (China). Antibodies to anti-V5-Tag (ab9116) and Anti-HA-Tag (ab9110) for immunoprecipitation were obtained from Abcam. Antibodies to Anti-Flag-tag (F1804) for immunoprecipitation were obtained from SIGMA. Antibodies to Ub (ab7780), ILF3/(NF90) (ab92355), and PKR (ab184257) were obtained from Abcam. The antibody to TRIM47 (BC017299) was obtained from Thermo (PA5-50892), and the antibody to Tim-3 (A2516) for western Blot was obtained from Abclonal.

### Western Blot Analysis

Western blots were performed as described previously(*36*). Briefly, cells were lysed with lysis buffer (1% Triton X 100, 20 mM Tris-HCl pH 8.0, 250 mM NaCl, 3 mM EDTA pH 8.0), 3 mM EGTA (pH 8.0) with the pH adjusted to 7.6, and complete protease inhibitor cocktail (Roche, pH 7.5) on ice for 30 min. Lysates were eluted by boiling 10 min with 5 Χ sample buffer (100 mM Tris-HCl, pH 6.8, 2% SDS, 10% glycerol, 0.1% bromophenol blue, 1% β-mercaptoethanol) and were separated by 10% SDS/PAGE, followed by examination of expression levels of the indicated proteins: phospho-eIF2α, eIF2α, phospho-PKR, PKR, total protein of NF90, and the levels of phospho-ERK, and phospho-p38. β-Actin served as an internal control.

### Co-Immunoprecipitation

Cells were collected 24 h after transfection and lysed in lysis buffer (1% NP-40, 20 mM Tris-HCl, 150 mM NaCl, 5mM EDTA, 1mM Na3VO4, 0.25% sodium deoxycholic acid and complete protease inhibitor cocktail (Roche), pH 7.5) on ice for 30 min. After centrifugation for 15 min at 12 000 r(11800 × *g*), 4°C, the supernatants were collected and incubated with Protein A / G Sepharose beads (SC-2003, Santa Cruz) coupled to specific antibodies overnight at 4 °C. The next day, beads were washed three times with high salt wash buffer (1% Triton × 100, 20 mM Tris-HCl, 500 mM NaCl, 10% Glycerol, 2 mM EDTA, 1 mM Na3VO4, and complete protease inhibitor cocktail (Roche), pH 7.5) and three times with low salt wash buffer (1% Triton × 100, 20 mM Tris-HCl, 150 mM NaCl, 10% Glycerol, 2 mM EDTA, 1 mM Na3VO4, and complete protease inhibitor cocktail (Roche), pH 7.5), respectively. Lysates were eluted by boiling 10 min with 5X sample buffer (as indicated). Precipitates were fractionated by SDS/PAGE with appropriate concentration and western Blot was performed as described above.

### Ubiquitination assays

For analysis of the ubiquitination of NF90 in HEK293T cells, plasmids encoding Flag-NF90, HA-Ub-K48 were transfected into HEK293T cells for 24 h and were treated with MG132 (20 μM) for 6 h before harvesting. Cells were lysed with IP lysis buffer (1% NP-40, 20 mM Tris-HCl, 150 mM NaCl, 5 mM EDTA, 1 mM Na3VO4, 0.5% sodium deoxycholic acid, and complete protease inhibitor cocktail (Roche), pH 7.5), and then the whole-cell lysates were immunoprecipitated with an antibody to Flag tag (F1804), followed by analysis of ubiquitination of NF90 with an antibody to HA tag. Precipitates were fractionated by SDS/PAGE with appropriate concentration (as indicated).

### Pathology and Survival assays

Survival of ∼6-week-old wildtype (*Tim-3*^*+/+*^) and Tim-3 knock out (*Tim-3*^−/−^) mice were given intraperitoneal injection with VSV (1 × 10^8^ pfu/g) (n = 9 per group). To detect the pathology of *Tim-3*^*+/+*^ and *Tim-3*^−/−^ mice in response to VSV, the hematoxylin and eosin staining of lung sections were examined 24 h after infecting.

### Mass spectrometry

Plasmid encoding full-length Tim-3 were transfected into HEK293T cells for 24 h, and cell lysates were immunoprecipitated with an antibody to Tim-3 (D5D5R #45208). Mass spectrometry was used to identify Tim-3-interacting proteins.

### Q-PCR and RNAi knockdown

Gene expression was analyzed by three-step q-RT–PCR (qPCR). Total RNA was extracted from mouse macrophages using TRI reagent (Sigma). Following the manufacturer’s instructions RNA was reverse-transcribed in a 20 μl reaction volume (42°C, 30 min; 95°C, 5 min) using a QuantiTect Reverse Transcription Kit (Qiagen, Valencia, CA, USA). cDNA was then amplified using a SYBR Green I Master mix (Roche, Basel, Switzerland) and a Light Cycler 480 PCR system (Roche). All tests were carried out on duplicate reaction mixtures in 96-well plates. The relative expression of the gene of interest was determined using the 2^−ΔΔCt^ method, with 18S ribosomal mRNA (18S) as the internal control. The primers used for qPCR are listed in Supplemental Figure 1.

### Statistical Analysis

The significance of difference between groups was determined by two-tailed Student’s t-test and two-way analysis of variance test. For the mouse survival study, Kaplan–Meier survival curves were generated and analyzed for statistical significance with GraphPad Prism 6.0. *P*-values <0.05 were considered statistically significant.

## Acknowledgments

We thank Prof. Minghong Jiang, Institute of Basic Medical Sciences, Peking Union Medical College, Chinese Academy of Medical Sciences, Beijing, China, for critical reading.

## Author Contributions

**Conceptualization:** Gencheng Han

**Data Curation:** Gencheng Han; Shuaijie Dou

**Formal Analysis:** Renxi Wang; Yang Zheng

**Investigation:** Shuaijie Dou; Ge Li

**Methodology:** Shuaijie Dou; Guoxian Li

**Project Administration:** Gencheng Han

**Resources:** Lili Tang; Yang Gao; Rongliang Mo; Beifen Shen;

**Software:** Ge Li; Chunmei Hou, Yuxiang Li;

**Supervision:** Chunmei Hou

**Validation:** Guoxian Li; Ge Li;

**Visualization:** Jun Zhang

**Writing – Original Draft Preparation:** Shuaijie Dou; Jun Zhang

**Writing – Review & Editing:** Gencheng Han

## Abbreviations

NF90: Nuclear Factor 90
SGs: Stress granules
VSV: Vesicular Stomatitis Virus
PKR: Protein kinase R
(eIF2α): the eukaryotic translation initiation factor 2α
G3BP1: Ras-GAP SH3-binding protein-1
TIA-1: T-cell intracellular antigen-1
TRIM47: tripartite motif-containing protein 47

